# Phosphatase to kinase switch of a critical enzyme contributes to timing of cell differentiation

**DOI:** 10.1101/2023.04.11.536480

**Authors:** Trisha N. Chong, Saumya Saurabh, Mayura Panjalingam, Lucy Shapiro

**Author notes:** Correspondence (L.S.), (S.S.).

## Abstract

Cell differentiation is an essential biological process that is often subject to strict temporal regulation. The aquatic bacterium, *Caulobacter crescentus*, undergoes obligate differentiation from a swarmer cell to a stationary, replication-competent stalked cell, with each cell cycle. Here, we report that the switch from phosphatase to kinase activity of the histidine kinase PleC contributes to timing this differentiation event. We show that PleC PAS domain interaction with the polar scaffold protein PodJ localizes PleC to the cell pole and inhibits *in vivo* kinase activity. Upon PodJ degradation, released PleC switches to its kinase form and phosphorylates the PleD diguanylate cyclase, initiating the signaling pathway responsible for differentiation. While PodJ inhibits PleC kinase activity, it does not impact PleC phosphatase activity on DivK, which is required for pili biogenesis and flagellar rotation. Thus, PleC PAS domain interaction with PodJ regulates PleC subcellular localization, enzymatic activity, and the timing of cell differentiation, revealing that PAS domains affect enzymatic function on diverse substrates by relying on context dependent binding partners.

## Introduction

Cell differentiation is the process by which a cell acquires a new cell fate that makes it specifically suited for its environmental or developmental context. The process of cell differentiation can be triggered by either an external cue or an internal signaling circuit. The aquatic bacterium, *Caulobacter crescentus*, has a di-morphic cell cycle in which a swarmer cell differentiates into a sessile, replication-competent stalked cell in response to both external and internal regulatory pathways (Figure 1A).

**Figure 1.**
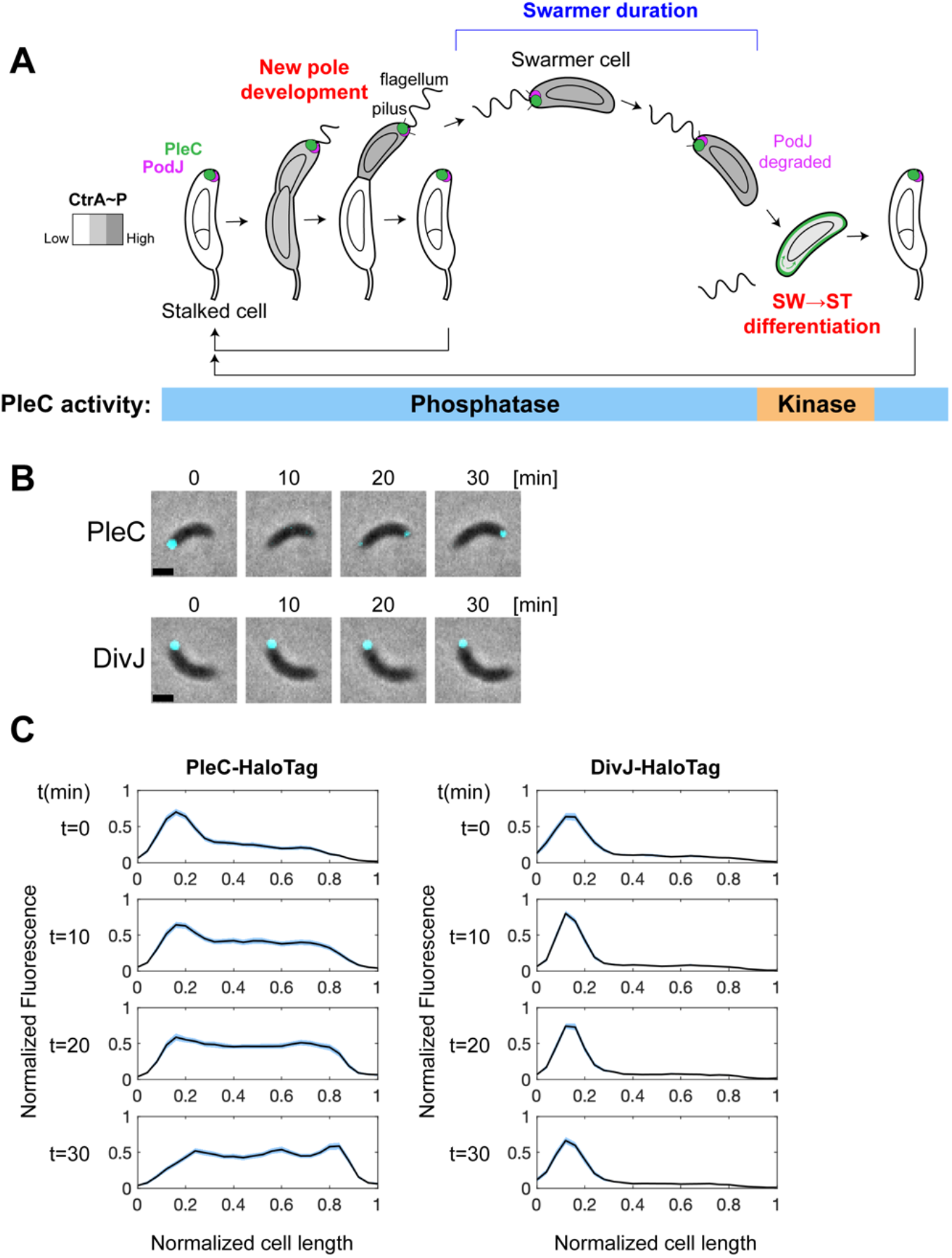
PleC molecules exhibit polar translocation during differentiation. **(A)** Schematic of PleC, PodJ and CtrA∼P localization throughout the cell cycle and the corresponding PleC enzymatic state resulting from PodJ inhibition of PleC kinase activity at the cell pole. **(B)** Representative merged phase and fluorescence time-lapse images of PleC-HaloTag (top) and DivJ-HaloTag (bottom) molecules pulse labeled with 2nM JF549 dye for 10 minutes. For PleC-HaloTag, cells were labeled immediately after synchrony. For DivJ-HaloTag, swarmer cells were allowed to progress to stalked cells for 40 minutes prior to labeling. Scale bar = 1μm. **(C)** Normalized fluorescence intensity profiles along normalized cell length axes of PleC-HaloTag (n=47) and DivJ-HaloTag (n=52) at ten-minute time points. Light blue shaded area denotes the standard error of the mean.

Cell division in *Caulobacter* yields a motile swarmer cell and a stationary stalked cell which immediately initiates DNA replication. The swarmer daughter cell must differentiate into a stalked cell before undergoing replication. Swarmer to stalked cell differentiation involves shedding the polar flagellum, proteolysis of the chemotaxis machinery, pili retraction, stalk biogenesis, and DNA replication initiation. These developmental changes are made possible by a cascade of molecular signaling events which ultimately lead to the dephosphorylation and proteolysis of the CtrA master cell cycle regulator (Figure 3A)(Domian et al., 1997; Quon et al., 1998). Later in the cell cycle, when newly synthesized CtrA is activated by phosphorylation, CtrA∼P promotes the transcription of over 90 genes involved in the development of the new flagellar pole (Figure 1A and 6A)(Laub et al 2002). A critical component of this molecular signaling cascade is the histidine kinase, PleC, which acts as either a kinase or a phosphatase at different times in the cell cycle. To initiate cell differentiation, the PleC kinase phosphorylates the diguanylate cyclase, PleD, leading to an increase in the second messenger, cyclic-di-GMP (c-di-GMP) (Paul et al., 2004). C-di-GMP directly binds to a ClpXP protease adaptor protein, activating ClpXP mediated proteolysis of CtrA and enabling swarmer to stalked cell differentiation (Figure 3A)(Joshi et al., 2015). On the other hand, PleC phosphatase activity on the response regulator DivK in pre-divisional cells leads to downstream phosphorylation and activation of newly synthesized CtrA, which is required for the development of the new flagellar pole, including flagellar rotation and pili biogenesis (Figure 6A)(Matroule et al., 2004; Tsokos et al., 2011). Mutants lacking PleC phosphatase activity are immotile and unable to form pili. This phenotype can be rescued by restoring specifically PleC phosphatase and not kinase activity (Matroule et al., 2004).

While the signaling pathway that governs swarmer to stalked cell differentiation has been extensively characterized, the mechanisms responsible for timing differentiation are poorly understood. Multiple reports have shown that interaction with a surface can trigger swarmer to stalked cell differentiation, resulting in a shortened swarmer cell duration. This is facilitated by signal transduction through either the polar pili or the flagellum (Ellison et al., 2017; Hershey et al., 2021; Hug et al., 2017; Medico et al., 2020; Snyder et al., 2020). However, when grown in nutrient rich liquid media, isolated swarmer cells reliably undergo differentiation into stalked cells after approximately 40 minutes (Degnen & Newton, 1972). This implies that in the absence of surface stimulation, an internal molecular circuit controls the timing of cell differentiation. It has been proposed that allosteric binding of DivK to PleC at the incipient stalked pole switches PleC to its kinase conformation, enabling robust phosphorylation of PleD leading to swarmer to stalked cell differentiation (Paul et al., 2008).

Here, we provide evidence that PleC interaction with the polar scaffold protein PodJ recruits PleC to the cell pole and maintains PleC in its phosphatase conformation when the two proteins colocalize at the cell pole (Figure 1A)(Hinz et al., 2003; Lawler et al., 2006; Zhang et al., 2022). For a brief period, prior to cell differentiation, the cytoplasmic domain of PodJ is proteolyzed (Chen et al, 2006), releasing PleC from the cell pole and enabling its kinase activity. PleC kinase phosphorylates PleD, initiating the molecular pathway that results in swarmer to stalked cell differentiation (Figure 3A). After differentiation, diffuse PleC is captured by newly synthesized PodJ at the opposite pole, reverting PleC back to its phosphatase form. In this way, PodJ proteolysis enables a PleC phosphatase to kinase switch, and initiates the molecular pathway that results in swarmer to stalked cell differentiation. It has been shown *in vitro* that PodJ binds directly to PleC through PleC’s cytoplasmic PAS domains (Zhang et al., 2022). Here, we show that a mutant lacking PleC PAS domains, whose kinase activity is not inhibited by PodJ, prematurely degrades CtrA and initiates the early onset of chromosome replication. Notably, only PleC kinase activity on PleD and not its phosphatase activity on DivK is affected by PodJ inhibition. We demonstrate that PodJ interaction with PleC’s PAS domains coordinates PleC’s enzymatic activity with its subcellular localization *in vivo*. Consequently, PleC is a phosphatase while colocalized with PodJ at the cell pole and a kinase only when diffuse, resulting in the phosphorylation of PleD and the initiation of swarmer to stalked cell differentiation during a narrow window of the cell cycle (Figure 1A).

## Results

### PleC molecules exhibit polar translocation during differentiation

PleC localizes to the flagellar pole throughout most of the cell cycle except for a brief period during swarmer to stalked cell differentiation, when PleC is observed to vacate the old flagellar pole and appear at the opposite, incipient flagellar pole (Figure 1A)(Matroule et al., 2004; Wheeler & Shapiro, 1999). To determine whether PleC appearing at the incipient flagellar pole is comprised of only newly synthesized PleC molecules or those that are released from the old flagellar pole, we performed a pulse labeling experiment on cells with PleC C-terminally fused to HaloTag (Los et al., 2008) as the sole copy of PleC encoded at the endogenous *pleC* locus. We pulse labeled synchronized swarmer cells with the fluorescent dye, Janelia Fluor 549 (JF549), that covalently binds to HaloTag, and subsequently washed away excess unbound dye. This allowed fluorescent tagging of PleC-HaloTag molecules that were present at the start of the time-lapse experiment. We observed that pulse-labeled PleC-HaloTag molecules originating from the old flagellar pole appeared as a fluorescent focus at the new flagellar pole (Figure 1B and 1C). As a negative control, we observed that pulse-labeled DivJ-HaloTag remained at the stalked pole during stalked cell development into pre-divisional cells (Figure 1B and 1C). PleC translocation occurred at the same time in the cell cycle that PodJ is released from the membrane and fully degraded (Chen et al., 2006). As PodJ is required for PleC polar localization (Figure 2A)(Hinz et al., 2003), release of PleC from the old flagellar pole is likely due to PodJ degradation at that pole.

**Figure 2.**
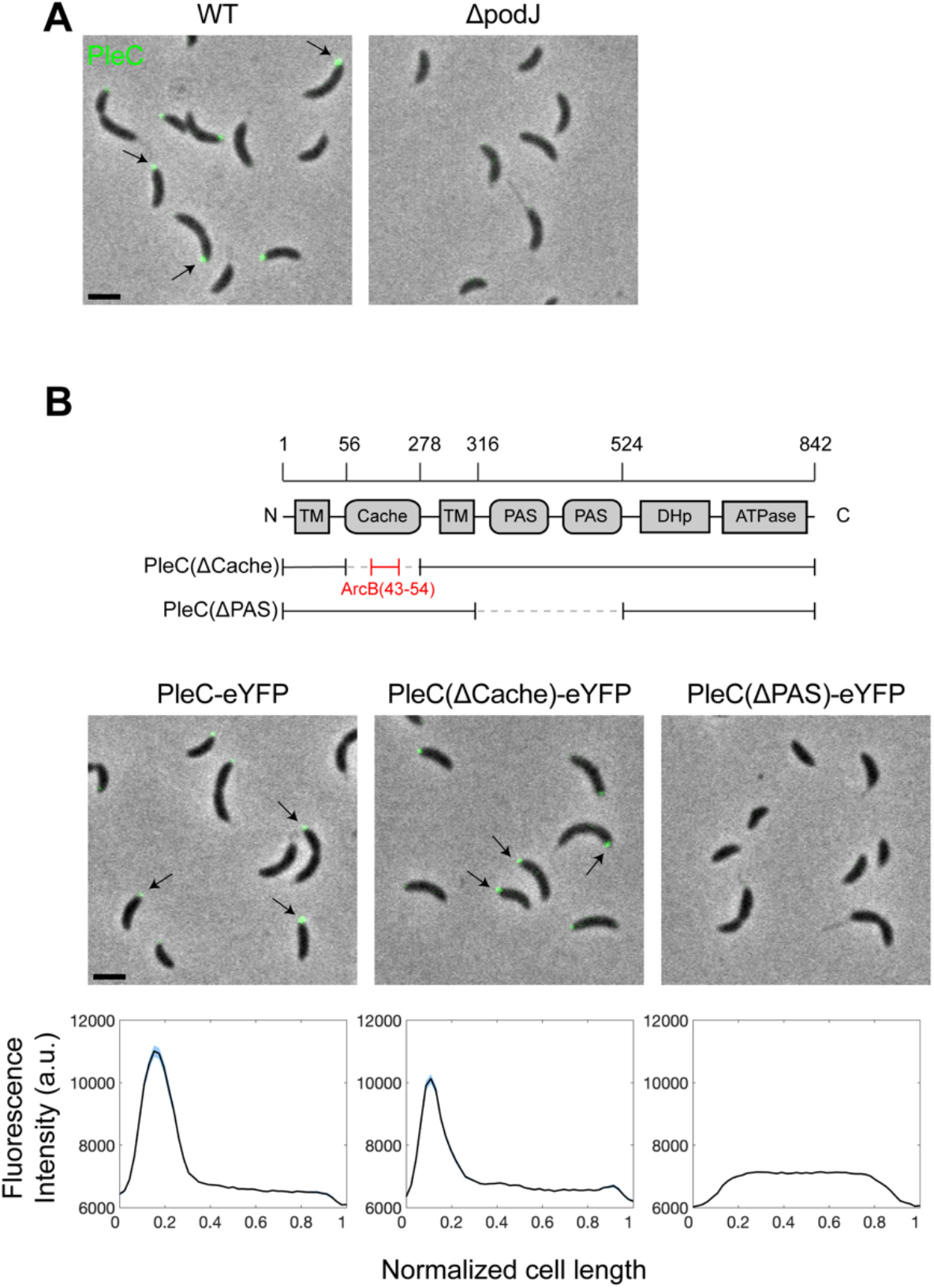
PleC polar localization is dependent on PAS domain interaction with PodJ. **(A)** Representative merged phase and fluorescence images of PleC C-terminally fused to eYFP in WT and *ΔpodJ* strains. Arrows point polar fluorescence signal. **(B)** Schematic of PleC domain architecture and PleC domain mutants (top). The *pleC(ΔCache)* variant was generated by replacing the periplasmic Cache domain of PleC with the unstructured periplasmic sequence from the *E. coli* protein ArcB. Representative merged phase and fluorescence images (middle) of PleC domain mutants with C-terminal eYFP. Arrows point to polar fluorescence signal. Scale bar = 2μm. Line profiles of average fluorescence intensity along normalized cell lengths (bottom). For each mutant, 1,189-1,415 cells were analyzed. Light blue shaded area denotes standard error of the mean.

We observed a reduction in PleC-HaloTag signal when comparing the intensity of the fluorescent focus at the old flagellar pole to the new flagellar pole (Figure 1B and 1C). This reduction in signal was greater than that expected by photobleaching, suggesting that not all PleC-HaloTag molecules translocated to the new pole. In addition to PleC polar translocation, it has been reported that some PleC proteolysis occurs at this point in the cell cycle (Coppine et al., 2020). Additionally, as PleC is synthesized throughout the cell cycle (Schrader et al., 2016; Zhou et al., 2015), newly synthesized PleC-HaloTag that was not labeled with JF549 likely localized to the new pole, diluting the fluorescent signal. Together these results suggest that during differentiation, PleC is released from the old flagellar pole upon PodJ proteolysis and then captured at the incipient flagellar pole by newly synthesized PodJ localized at that pole (Figure 1A, 1B and 1C).

### PleC polar localization is dependent on PAS domain interaction with PodJ

PleC polar localization is dependent on the polar scaffold protein, PodJ (Figure 2A)(Hinz et al., 2003). PleC is a multi-domain histidine kinase (Figure 2B)(Zhang et al., 2022). The N-terminus of PleC consists of a periplasmic Cache domain positioned between two transmembrane domains. Cache domains are extracellular or periplasmic sensory domains that are homologous to, but structurally distinct from, PAS domains (Upadhyay et al., 2016). In the cytoplasm, PleC contains two PAS (<u>P</u>er-<u>A</u>rnt-<u>S</u>im) domains, a DHp (<u>D</u>imerization and <u>H</u>istidine <u>p</u>hosphotransfer) domain and a C-terminal ATPase domain (Figure 2B)(Zhang et al., 2022). To identify the domains that are required for PleC polar localization, we constructed two domain deletion mutants: a Cache domain deletion and a PAS domain deletion, both C-terminally fused to enhanced yellow fluorescent protein (eYFP) and expressed as the sole copy of PleC from its native promoter (Figure 2B). The *pleC(ιCache)* mutant was constructed by replacing the periplasmic Cache domain with the periplasmic linker portion of the *E. coli* protein ArcB, as this protein should not interact with any proteins specific to *Caulobacter* (Figure 2B).

WT PleC fused to eYFP localized to the flagellar pole of swarmer cells and to the incipient flagellar pole of stalked and pre-divisional cells (Figure 2A and Figure 2B)(Wheeler & Shapiro, 1999). We confirmed that PleC-eYFP is not polarly localized in **Δ*podJ* cells, although PleC is detectable at WT levels by Western blot (Figure 2A and Figure S1). We observed that the average intensity of the fluorescent polar focus of *pleC(ΔCache)-eYFP* cells was approximately 30% lower compared to WT cells (Figure 2B). In contrast, PleC(ιPAS)-eYFP was diffuse in swarmer, stalk, and pre-divisional cells (Figure 2B). We confirmed the stable expression of PleC(*Δ*Cache) and PleC(*Δ*PAS) by Western blot (Figure S1). It has been reported that PleC PAS domains bind directly to the cytoplasmic portion of PodJ *in vitro* (Zhang et al., 2022). Therefore, PleC(*Δ*PAS)-eYFP is likely diffuse because it is not able to bind to and colocalize with PodJ at the cell pole.

### PleC kinase activity is inhibited by PodJ interaction with PleC PAS domains

The PleC kinase phosphorylates the receiver domain of the diguanylate cyclase PleD (Figure 3A)(Paul et al., 2007, 2008). To investigate the effect of PodJ binding to PleC on its kinase activity *in vivo*, we performed PhosTag gel-electrophoresis to measure the ratio of PleD∼P/PleD in cells expressing PleD-HaloTag-3xFlag as the sole copy of PleD expressed from the endogenous locus (Barbieri & Stock, 2008; Kinoshita et al., 2006). We found that the ratio of PleD∼P/PleD was higher in *pleC(ΔPAS)* (155±28% of WT) compared to both WT and *ΔpleC*, suggesting that deleting the PAS domains results in increased kinase activity on PleD (Figure 3B). This result agrees with previous observations that PodJ inhibits PleC autokinase activity *in vitro* (Zhang et al., 2022). Because the ratio of PleD∼P/PleD in *ΔpleC* (83±35% of WT) was not significantly lower than that of WT, we suspect that PleC might act as both a kinase and a phosphatase on PleD and that this activity is cell cycle dependent (Figure 3B). Together these results suggest that direct binding of PodJ to PleC PAS domains both enables PleC polar localization and inhibits PleC kinase activity on PleD (Figure 2A, 2B and 3B).

When phosphorylated, PleD synthesizes the second messenger, c-di-GMP (Figure 3A)(Paul et al., 2004). C-di-GMP has been shown to have a role in regulating the lifestyle switch from planktonic to biofilm forming cells in many bacteria including *E. coli, Pseudomonas aeruginosa, Salmonella enterica* serovar Typhimurium, and others (Valentini & Filloux, 2016). In *Caulobacter*, c-di-GMP has been previously implicated in reducing motility and stimulating swarmer to stalked cell differentiation (Abel et al., 2013). To test whether the increased PleD∼P/PleD ratio in *pleC(ΔPAS)* coincides with reduced swarm ability, we seeded colonies onto semi-solid peptone yeast extract (PYE) plates and observed the increase in swarm area over time. As expected, *Δ*pleC colonies expanded to an area much smaller (10±1% of WT) than that of WT (Figure 3C)(Burton et al., 1997). In contrast, the swarm area of *pleC(ΔPAS)* (50±2% of WT) was smaller than that of WT but not as small as that of *ΔpleC* (Figure 3C). We also observed that the swarm area of *ΔpodJ* was smaller than that of *pleC(ΔPAS)*, suggesting that the *podJ* deletion has other pleiotropic effects related to swarming that are independent of PodJ’s interaction with PleC (Figure 3C). These pleiotropic effects might be due to insufficient PleD synthesis in *ΔpodJ* (Figure S2)(Guzzo et al., 2021). As swarm areas measure the average motility and chemotaxis of a population of cells over the course of a few days, we reasoned that the increased kinase activity of PleC(11PAS) on PleD could lead to decreased swarm area through two possible mechanisms: (1) reduced speed of swarmer cells (Nesper et al., 2017), or (2) premature swarmer to stalked cell differentiation, shortening the duration of the swarmer phase of the cell cycle.

**Figure 3.**
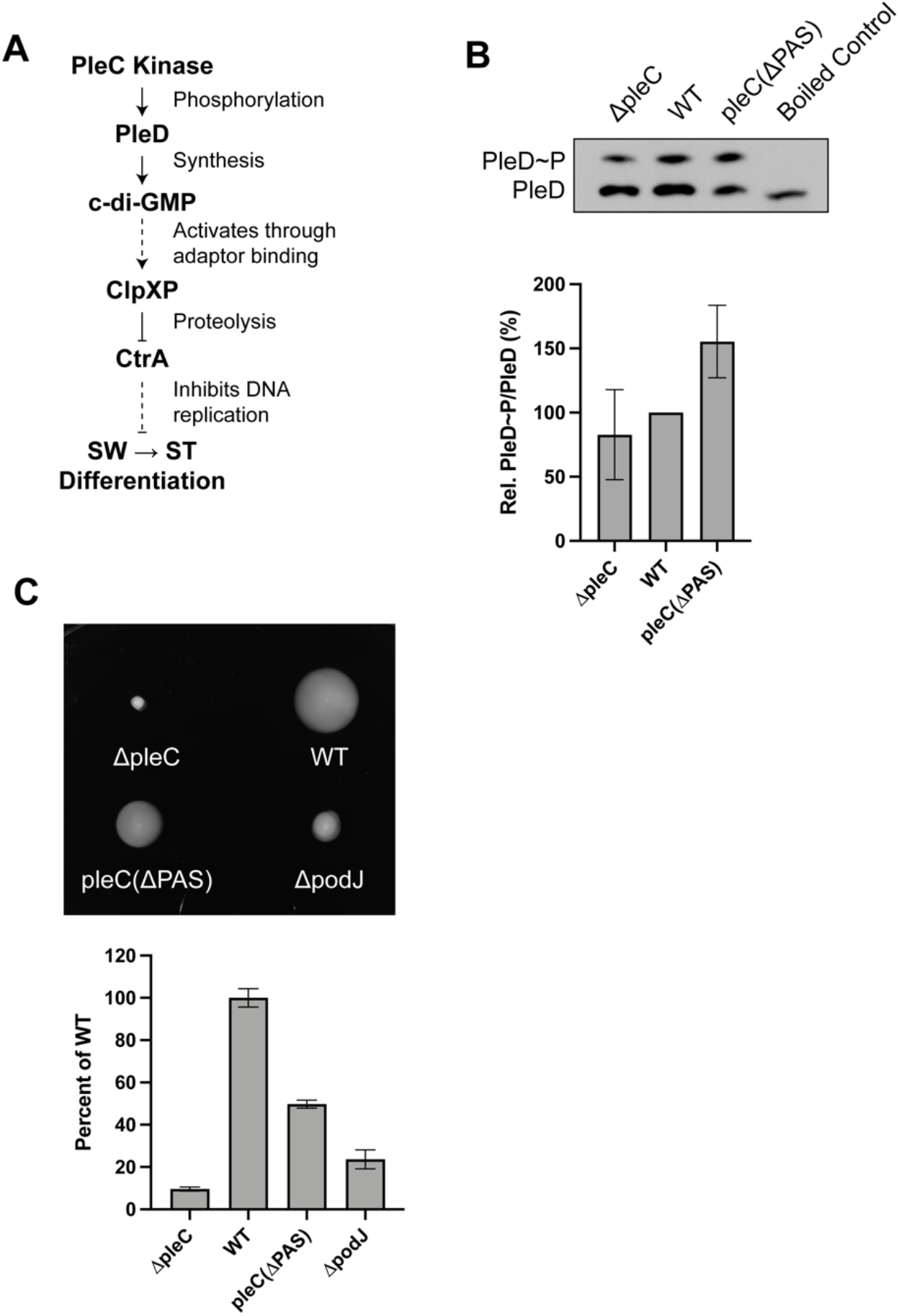
PleC kinase activity is inhibited by PodJ interaction with PleC PAS domains **(A)** The signal transduction pathway from PleC kinase activity on the diguanylate cyclase, PleD, that leads to swarmer (SW) to stalked (ST) cell differentiation. **(B)** PhosTag SDS-PAGE immunoblots showing phosphorylated and unphosphorylated PleD-HaloTag-3xFlag using an anti-Flag antibody (top). Bar plot shows average ratios of phosphorylated to unphosphorylated protein as a percentage of the WT ratio (bottom). Signal density was averaged across four replicates. Error bars indicate standard deviation. **(C)** Swarm plates showing the swarm abilities of mutants in PYE 0.3% agar over three days (top). Bar plot shows swarm areas as a percentage of WT averaged across ten replicate plates (bottom). Error bars indicate standard deviation.

### Loss of PAS domains leads to hypermotility

Swarm assays represent a population average of motility and chemotaxis effects. To overcome the limitations of the swarm assay, we quantified motility at the single cell level. Using phase contrast microscopy, we observed the swimming behavior of individual swarmer cells in glass bottom wells. We observed cells at a focal plane above the coverslip as to not capture cells that had settled to the bottom of the well. The majority of WT and *pleC(ΔPAS)* swarmer cells were observed swimming in straight or curved lines, characteristic of *Caulobacter’s* run and flick motility (Figure 4A)(Morse et al., 2016). As a negative control, we observed the flagellar mutant, *ΔfliG*, since unlike most *Caulobacter* strains, *ΔpleC* swarmer cells cannot be isolated by density centrifugation (Ardissone et al., 2014; Ramakrishnan et al., 1994). In contrast to WT and *pleC(ΔPAS)* swarmer cells, the majority of *ΔfliG* cells exhibited small twitching movements, even when observed away from the coverslip (Figure 4A). To gain further insights into the swimming behavior of these cells, we performed segmentation and tracking using the Fiji plugin, TrackMate (Tinevez et al., 2017). These analyses allowed for mapping the trajectories of individual swarmer cells and quantification of their instantaneous speeds along those trajectories (Figure 4A and 4B). To observe the extent of motility of each strain, we set the starting point of each trajectory to the origin. This analysis showed that *pleC(ΔPAS)* swarmer cells would cover an area similar to that of WT over the same amount of time (Figure 4A). In addition, the distribution of instantaneous speeds showed that both WT and *pleC(ΔPAS)* had higher fractions of cells that swam at fast instantaneous speeds (>30μm/s) compared to *ΔfliG* (Figure 4B). Interestingly, we discovered that a slightly higher fraction of *pleC(ΔPAS)* cells swam at fast instantaneous speeds (>30μm/s) compared to WT (Figure 4B). This analysis led us to conclude that the difference in swarm areas between WT and *pleC(ΔPAS)* is not due to differences in the speed of swarmer cells between the two strains. We next considered that a shortened swarmer duration of the cell cycle could account for the reduced swarm size of *pleC(ΔPAS)*.

**Figure 4.**
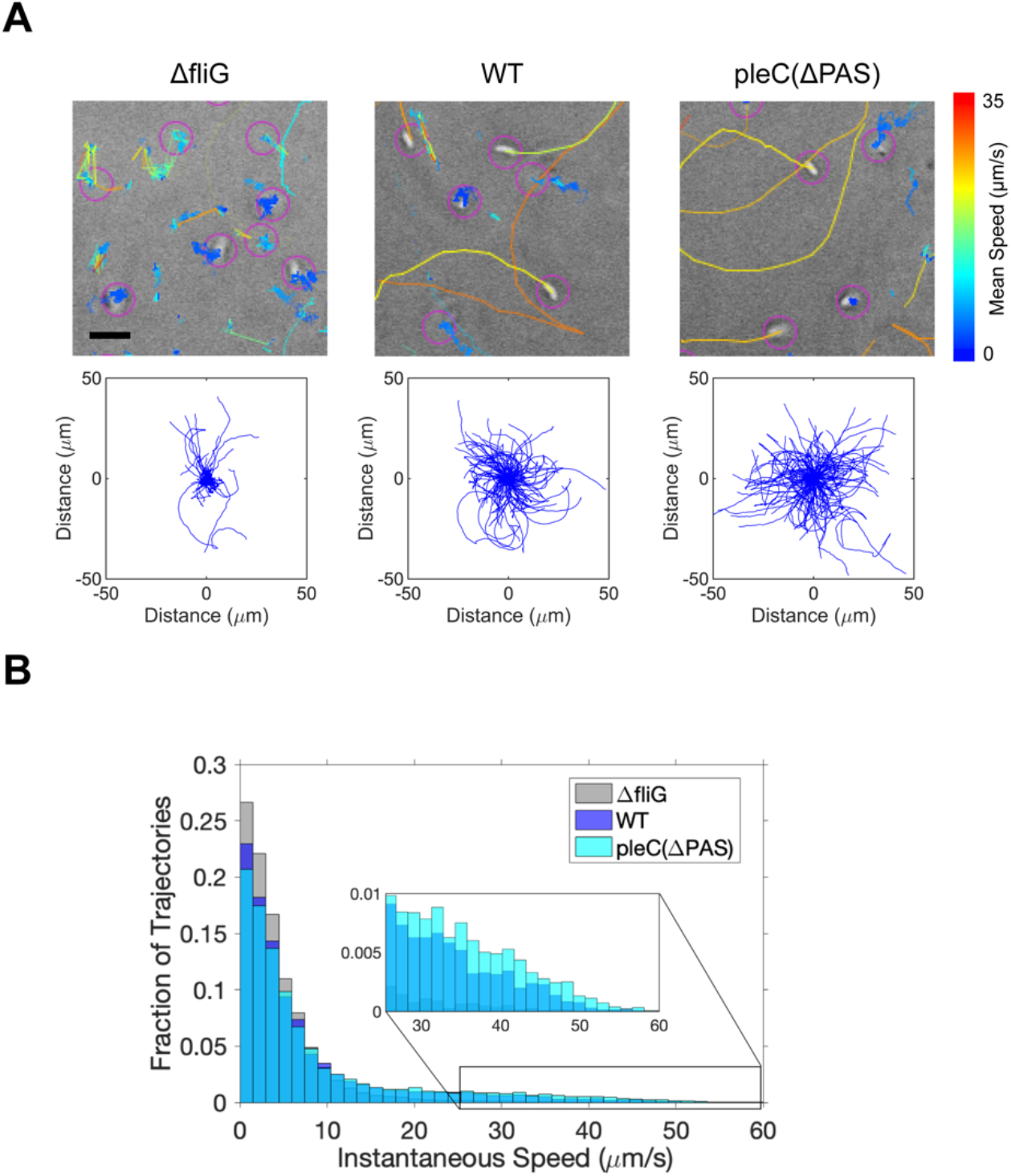
Loss of PAS domains leads to hypermotility. **(A)** Representative tracks of individual swarmer cells in M2G with 20% glycerol taken over a time interval of 10.4s (top). Track color corresponds to the track’s mean speed. Scale bar = 4μm. Plots of trajectories taken over a time interval of 26s with starting time points mapped to coordinate (0,0) (bottom). For each mutant, 806-1,625 trajectories were analyzed. **(B)** Histogram of the fraction of total trajectories in bins corresponding to instantaneous speed values. Zoomed in portion of the histogram shows that a larger fraction of *pleC(ΔPAS)* cells have high instantaneous speed values compared to WT cells.

### Loss of PleC PAS domain interaction with PodJ results in premature swarmer to stalked cell differentiation

Multiple events occur during swarmer to stalked cell differentiation, including dephosphorylation and degradation of the master cell cycle regulator CtrA that enables the initiation of chromosome replication, degradation of the polar chemotaxis machinery, and remodeling of the new stalked pole (Bowman et al., 2010; Joshi et al., 2015; Quon et al., 1998; Tsai & Alley, 2001). To address the possibility that the reduced swarm size of *pleC(ΔPAS)* is due to a shortened swarmer phase of the cell cycle, we used these developmental events as cell cycle landmarks to determine the difference in swarmer duration between WT and *pleC(ΔPAS)*.

Western blots of synchronized cells showed that CtrA was degraded more rapidly in *pleC(ΔPAS)* and *ΔpodJ* mutant cells than in WT cells (Figure 5A and 5B). This is likely due to increased levels of c-di-GMP activating the CtrA protease, ClpXP (Figure 3A)(Joshi et al., 2015). In addition to CtrA, the chemotaxis machinery positioned at the flagellar pole is also degraded upon swarmer to stalked cell differentiation (Tsai & Alley, 2001). Western blots of synchronized cells showed that the chemoreceptor, McpA, was degraded more rapidly in *pleC(ΔPAS)* and *ΔpodJ* cells compared to WT, suggesting that not only the timing of CtrA proteolysis, but also degradation of the chemotaxis machinery is affected by the loss of PleC interaction with PodJ (Figure 5A). As a control, we probed the outer membrane protein, PAL, that does not change in abundance throughout the cell cycle (Figure 5A)(Yeh et al., 2010). These results suggest that swarmer cell-specific proteins are degraded more rapidly in *pleC(ΔPAS)* and *ΔpodJ* cells, suggesting that these mutants have shorter swarmer durations compared to WT.

**Figure 5.**
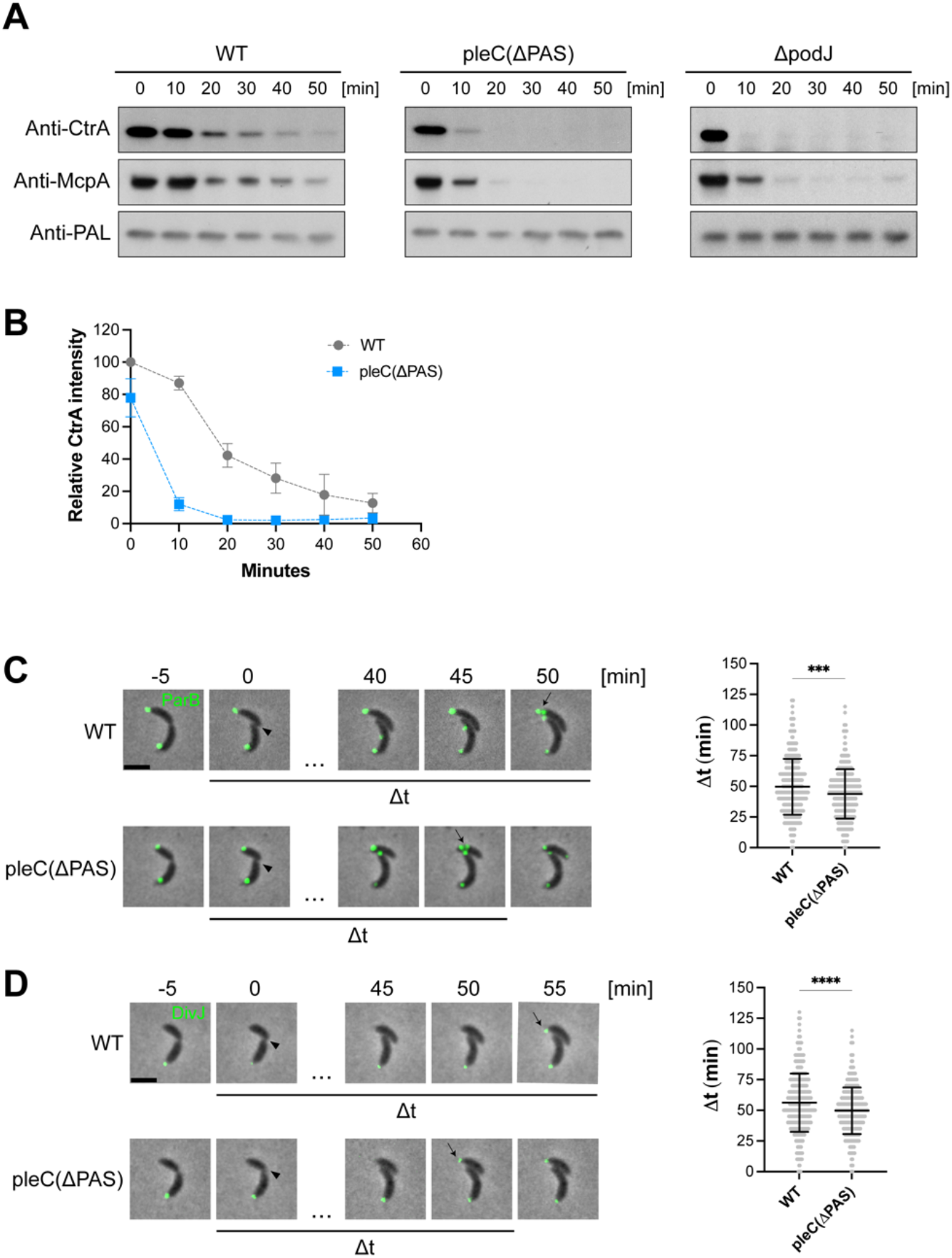
Loss of PleC PAS domain interaction with PodJ results in premature swarmer to stalked cell differentiation. **(A)** Western blot showing CtrA and McpA levels of synchronized WT, *pleC(ΔPAS)* and *ΔpodJ* cells. Western blot against non-cell-cycle-controlled protein PAL shows consistent levels through the cell cycle. **(B)** Quantification of CtrA signal density averaged across four replicates. WT and *pleC(ΔPAS)* samples were run on the same gel for each replicate. Error bars denote standard deviation. **(C)** Time-lapse images of eYFP-ParB foci as swarmer cells differentiate into stalked cells and initiate DNA replication. Representative merged phase and fluorescence time-lapse images of WT and *pleC(ΔPAS)* cells expressing *pxyl:eYFP-parB*, induced with 0.01% for 1 hour. Arrowheads point to cytokinesis events. Arrows point to a single eYFP-parB focus splitting into two foci. Scale bar = 2μm. Violin plot of Δt, the time interval between cytokinesis and splitting of the eYFP-parB focus into two foci (right). WT n = 390. *pleC(ΔPAS)* n = 362. Error bars denote the mean ± standard deviation. An unpaired t-test was used to calculate p=0.0002. **(D)** Time-lapse images of DivJ-eYFP foci as swarmer cells differentiate into stalked cells. Representative merged phase and fluorescence time-lapse images of WT and *pleC(ΔPAS)* cells with DivJ-eYFP (left). Arrowheads point to cytokinesis events. Arrows point to appearance of a new DivJ-eYFP focus. Scale bar = 2μm. Violin plot of Δt, the time interval between cytokinesis and appearance of a new DivJ-eYFP focus (right). WT n = 413. *pleC(ΔPAS)* n = 471. Error bars denote mean ± standard deviation. An unpaired t-test was used to calculate p<0.0001.

In a second assay to determine the timing of the swarmer to stalked cell differentiation, we measured the time between cytokinesis and initiation of chromosome replication. The ParB protein binds to the *parS* DNA sequence near the origin of replication, the first region of the chromosome that upon duplication is transported rapidly to the opposite cell pole (Toro et al., 2008). At the start of replication, the ParB protein bound to *parS* is located at the incipient stalked pole. Splitting of a single ParB focus into two distinct foci indicates duplication of the *parS* DNA sequence and the initiation of chromosome replication (Toro et al., 2008). Because initiation of chromosome replication occurs exclusively in stalked cells (Quon et al., 1998), this event can be used as a cell cycle landmark. To measure the duration of the swarmer phase of the cell cycle, we synchronized WT and *pleC(ΔPAS)* cells harboring *pxyl:eYFP-parB* and allowed swarmer cells to progress to pre-divisional cells prior to capturing time-lapse images. We measured the duration of the swarmer phase of the cell cycle beginning at cytokinesis and ending at the splitting of the single eYFP-ParB focus into two distinct foci (Figure 5C). We found that this time interval was significantly shorter for *pleC(ΔPAS)* compared to WT, with an average time difference of 5.9±1.6 minutes (Figure 5C). These results suggest that initiation of chromosome replication occurs after a shorter swarmer cell duration in *pleC(ΔPAS)* cells compared to WT cells.

In a third assay, we measured the time difference between cytokinesis and the appearance of the DivJ histidine kinase which localizes to the stalked pole (Wheeler & Shapiro, 1999). During swarmer to stalked cell differentiation, swarmer pole proteins are degraded and newly synthesized stalked pole proteins, including DivJ, localize to the new stalked pole (Bowman et al., 2010; Joshi et al., 2015). Accordingly, we measured the time interval between cytokinesis and the appearance of a DivJ-eYFP focus at the newly formed stalked pole. We found that this time interval was significantly shorter for *pleC(ΔPAS)* compared to WT with an average difference of 6.6±1.4 minutes (Figure 5D). Together these results suggest that *pleC(ΔPAS)* swarmer cells progress to stalked cells faster than WT swarmer cells and that PleC PAS domain interaction with PodJ plays a critical role in regulating the duration of the swarmer phase of the cell cycle.

### PleC polar localization is not required for DivK dephosphorylation and pili biogenesis

PleC phosphatase activity on DivK is required for downstream phosphorylation and activation of CtrA at the flagellar pole of pre-divisional cells (Figure 6A). The pseudokinase DivL directly binds to and promotes CckA autokinase activity in the absence of DivK∼P (Mann & Shapiro, 2018; Tsokos et al., 2011). When DivK is phosphorylated, DivL and DivK∼P form a complex that inhibits CckA kinase activity. Therefore, PleC phosphatase activity on DivK frees CckA from DivL-DivK∼P inhibition and allows CckA kinase activity and phosphorylation of CtrA by phosphotransfer through ChpT (Figure 6A)(Childers et al., 2014; Mann & Shapiro, 2018; Tsokos et al., 2011). When phosphorylated and active, CtrA promotes the transcription of over 90 genes that are involved in development of the new flagellar pole, which includes flagellar rotation and pili biogenesis (Figure 6A)(Laub et al., 2002; Matroule et al., 2004). To determine whether PleC phosphatase activity on DivK is impaired in *pleC(ΔPAS)* mutants, in which PleC is diffuse (Figure 2B), we performed PhosTag gel-electrophoresis to measure the ratio of DivK∼P/DivK in cells expressing DivK-HaloTag-3xFlag as the sole copy of DivK from the endogenous locus. We found that the ratio of DivK∼P/DivK was very high in *ΔpleC* (284±33% of WT), as would be expected from removing a DivK phosphatase (Figure 6B). We found that the ratio of DivK∼P/DivK in *pleC(ΔPAS)*(73±26% of WT) was similar to that of WT (Figure 6B). While we observed increased kinase activity on PleD in *pleC(ΔPAS)* compared to WT, we did not see a significant difference in phosphatase activity on DivK, suggesting that PleC in its kinase conformation retains some phosphatase activity (Figure 3B and 6B). Additionally, these results suggest that diffuse PleC(*Δ*PAS) does not have impaired phosphatase activity, and that PleC polar localization is not required for robust phosphatase activity on DivK (Figure 2B and 6B).

**Figure 6.**
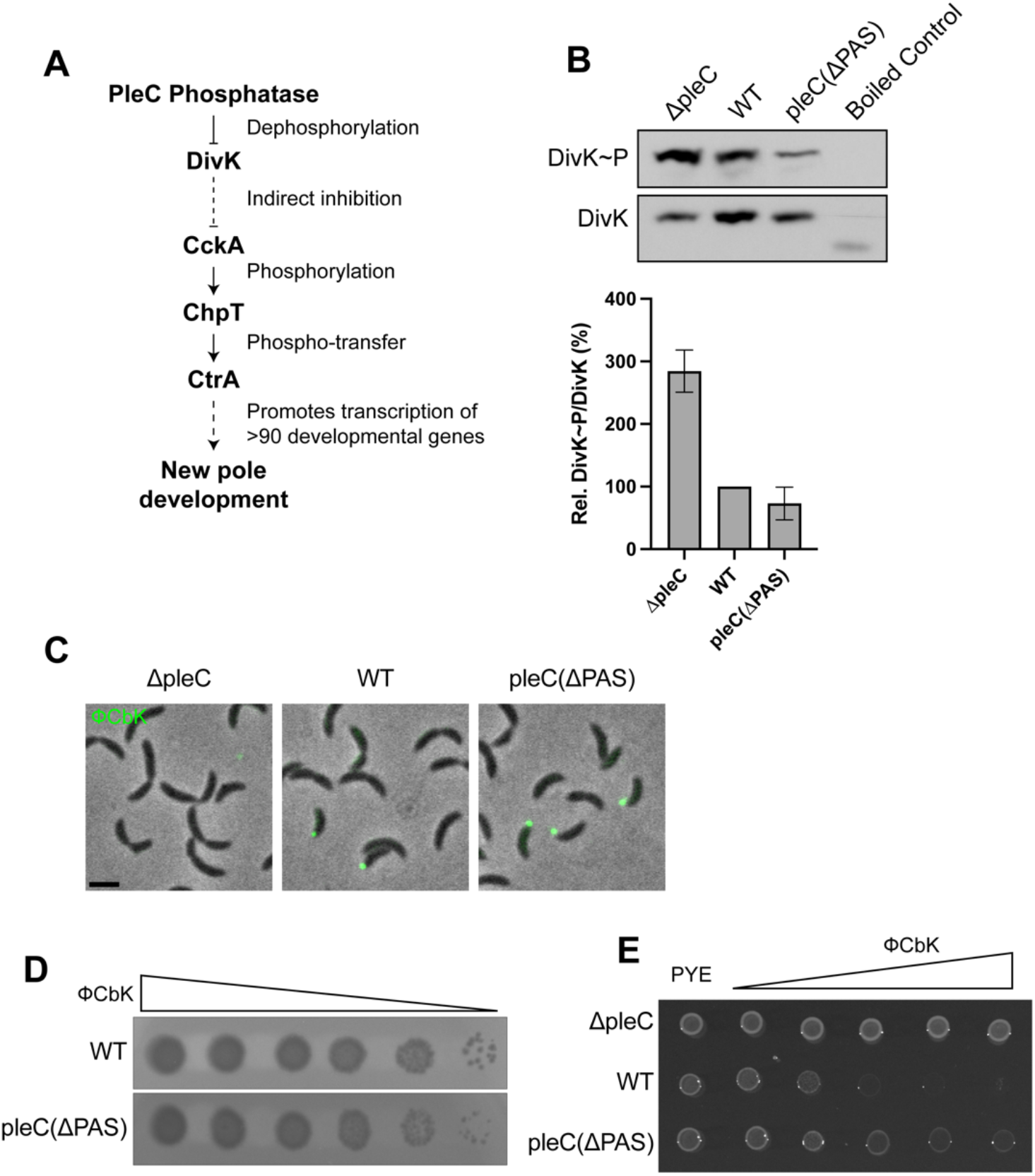
PleC polar localization is not required for DivK dephosphorylation and pili biogenesis. **(A)** Schematic of PleC phosphatase activity on DivK and the downstream molecular pathway that results in new pole development. **(B)** PhosTag SDS-PAGE immunoblots showing phosphorylated and unphosphorylated DivK-HaloTag-3xFlag using an anti-Flag antibody (top). Bar plot shows average ratios of phosphorylated to unphosphorylated protein as a percentage of the WT ratio (bottom). Separate exposure times were used to capture unsaturated phosphorylated and unphosphorylated bands. Signal density was averaged across two replicates. Error bars indicate standard deviation. **(C)** Fluorescent *ϕ*CbK phage labeled overnight with 25μM Sytox Green attaching to cells with polar pili. Scale bar = 2μm. **(D)** Serial dilutions of *ϕ*CbK spotted onto bacterial lawns that formed plaques overnight. **(E)** Cells grown overnight in the presence of serial dilutions of *ϕ*CbK.

It has been previously shown that PleC phosphatase and not kinase activity is required for development of the new flagellar pole, including flagellar rotation and pili biogenesis (Matroule et al., 2004). Because our results show that PodJ interaction with PleC PAS domains inhibits PleC kinase activity but does not impact PleC phosphatase activity, we suspected that new pole development would not be gravely impaired in *pleC(ΔPAS)* cells. To test this hypothesis, we assayed the ability of ϕCbK, a phage that infects *Caulobacter* through its polar pili, to attach to and infect *Caulobacter* cells (Guerrero-Ferreira et al., 2011; Skerker, 2000). We fluorescently labeled ϕCbK with Sytox green, a membrane impermeable DNA dye, and introduced labeled phage to *Caulobacter* cells. Phage that attached to pili appeared as fluorescent puncta at the cell pole. We observed fluorescent ϕCbK puncta at the cell poles of WT and *pleC(ΔPAS)* swarmer cells indicating that these cells have polar pili (Figure 6C). We did not see fluorescent ϕCbK puncta at the poles of *ΔpleC* cells which do not grow pili (Figure 6C)(Matroule et al., 2004). To assess whether cells were sensitive to phage infection, we spotted serial dilutions of ϕCbK onto bacterial lawns and observed plaque formation. Both WT and *pleC(ΔPAS)* cells formed plaques and are therefore susceptible to phage infection (Figure 6D). We observed that the plaques at the lowest phage concentration were bigger in WT than in *pleC(ΔPAS)*, suggesting that this mutant might have slightly higher resistance to ϕCbK compared to WT. When cells were incubated with ϕCbK, and plated on PYE plates, we observed that *pleC(ΔPAS)* cells showed an increased tolerance to high concentrations of phage compared to WT (Figure 6E). This increased tolerance to ϕCbK is likely due to only swarmer and late pre-divisional cells having pili and being susceptible to ϕCbK. Because *pleC(ΔPAS)* spends less time in the swarmer period of the cell cycle, fewer susceptible swarmer cells are present in a mixed population. Together these results suggest that PodJ interaction with PleC PAS domains does not impact PleC phosphatase activity and does not impair pili biogenesis at the new flagellar pole.

## Discussion

### PleC phosphatase to kinase switch initiates swarmer to stalked cell differentiation

PleC is a critical enzyme that functions as either a kinase or a phosphatase during distinct parts of the *Caulobacter* cell cycle. Here, we demonstrate *in vivo* that PodJ recruits PleC to the cell pole and inhibits PleC kinase activity on PleD through interaction with PleC’s PAS domains. PleC’s PAS domains have been shown *in vitro* to directly interact with the intrinsically disordered region (IDR) of PodJ, thereby inhibiting PleC autokinase activity (Zhang et al., 2022). Our data support a model in which release of PleC from the pole, upon PodJ degradation, enables phosphorylation and activation of PleD. At this time, concurrent allosteric binding of DivK to PleC might further stimulate PleC kinase activity on PleD (Paul et al., 2008). Phosphorylated PleD synthesizes c-di-GMP, which binds to the ClpXP adaptor, PopA, enabling the degradation of the master cell cycle regulator, CtrA (Figure 7A)(Joshi et al., 2015). Proteolysis of CtrA is required for initiation of chromosome replication which occurs exclusively in stalked cells (Quon et al., 1998). Increased c-di-GMP concentration also activates ShkA which initiates stalk pole development (Kaczmarczyk et al., 2020). Therefore, PleC release from the cell pole and phosphorylation of PleD initiates a cascade of signaling events that results in swarmer to stalked cell differentiation. After differentiation, newly synthesized PodJ localizes to the incipient flagellar pole, recruiting PleC to that pole, and reverting it back to its phosphatase form (Figure 1A and 7B).

**Figure 7.**
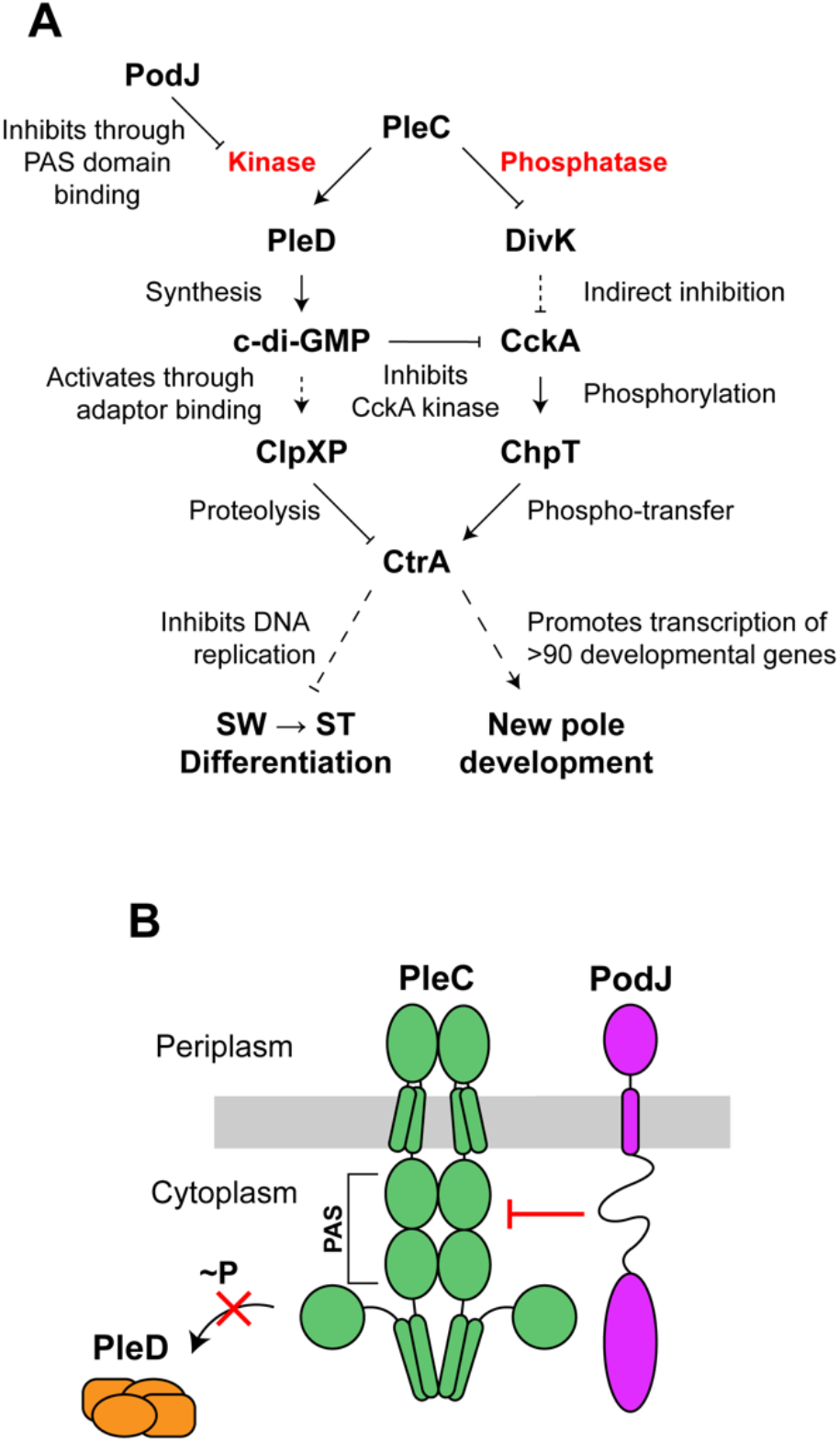
PodJ inhibition of PleC kinase activity controls timing of cell differentiation. **(A)** Schematic of molecular signaling pathways that result from PleC kinase and phosphatase activities and the downstream effects on swarmer (SW) to stalked (ST) cell differentiation and development of the new pole respectively. **(B)** Model of PodJ interaction with PleC PAS domains, resulting in inhibition of PleC kinase activity on PleD.

In prokaryotes, histidyl-aspartyl (His-Asp) signal transduction systems play a major role in cellular adaptation to environmental conditions. Most bacterial histidine kinases are bifunctional, having the ability to switch between kinase and phosphatase activity (Egger et al., 1997; Zhu et al., 2000). Some histidine kinases undergo conformational changes in response to environmental changes such as pH or osmolality, as in the cases of *Hermotoga maritima* HK853 or *Salmonella enterica* EnvZ (Kenney & Anand, 2020; Liu et al., 2017). Others change conformation through protein-protein interactions, such as PII-mediated phosphatase activation of NtrB (Weiss et al., 2002). In *Caulobacter*, the histidine kinase, CckA, switches between kinase and phosphatase activity depending on its subcellular localization and interaction partners at the cell pole (Mann et al, 2016). Through reconstitution of CckA on liposomes, it has been shown that autokinase activity is stimulated by high concentrations of CckA (Mann et al., 2016). Therefore, at the flagellar pole, where CckA concentration is high, CckA autokinase activity is stimulated in a density dependent manner. While the stalked pole also has a high CckA concentration, density-dependent autokinase activity is overshadowed by the inhibitory effects of direct binding of c-di-GMP and the DivL-DivK∼P complex to CckA, which have both been independently shown to inhibit CckA kinase and promote phosphatase activity (Lori et al., 2015; Mann et al., 2016; Mann & Shapiro, 2018; Tsokos et al., 2011). PleC is essential to the *Caulobacter* cell cycle, where it phosphorylates PleD and dephosphorylates DivK.

Here, we demonstrate that *in vivo*, PleC interaction with the scaffold protein PodJ at the cell pole inhibits PleC kinase activity on PleD, maintaining PleC in its phosphatase form during most of the cell cycle except for a narrow window of time in which PleC acts as a kinase following PodJ proteolysis (Figure 1A and 7B).

Previous work has shown that PleC primarily localizes at the flagellar pole, where it exclusively encounters DivK to dephosphorylate (Jacobs et al., 2001), as PleD is located at the stalked pole (Paul et al., 2004). PleC phosphatase activity on DivK is required for new pole development which includes flagellar rotation and pili biogenesis (Matroule et al., 2004). It has been proposed that PleC phosphatase activity at the flagellar pole ensures that DivK molecules at that pole are in the unphosphorylated state and unable to inhibit CckA kinase activity through direct interaction with DivL (Tsokos et al., 2011). CckA kinase activity at that pole leads to phosphorylation and activation of CtrA which promotes the transcription of genes that are required for new pole development (Figure 7A)(Mann et al., 2016; Matroule et al., 2004; Tsokos et al., 2011). Therefore, PleC kinase activity plays a critical role in timing the swarmer to stalked cell differentiation, while PleC phosphatase is required for new pole development prior to asymmetric cell division.

### Disruption of PodJ binding leads to PleC phosphatase to kinase switch

We showed that upon deletion of PleC’s PAS domains, PodJ was unable to inhibit PleC kinase activity, resulting in premature swarmer to stalked cell differentiation and a shortened swarmer duration of the cell cycle (Figure 3B, 5A, 5B, 5C and 5D). PodJ is sequentially degraded first by PerP cleavage of the PodJ periplasmic domain and then by MmpA which frees PodJ from the membrane, subsequently releasing PleC from the cell pole (Chen et al., 2005, 2006). In addition to PodJ proteolysis, other factors may contribute to PleC release from the cell pole and switch to its kinase form. Tan et al. showed that addition of purified SpmX protein to PodJ biomolecular condensates led to condensate disassembly and impaired recruitment of PodJ client proteins, including PleC (Tan et al., 2022). These results suggest that localization of newly synthesized SpmX to the incipient stalked pole might lead to dissociation of the PodJ polar complex and subsequent release of PleC from that pole. Medico et al. showed through bacterial-2-hybrid assays that PilA interacts with PleC’s N-terminal transmembrane domain and that *ΔpilA* swarmer cells take longer to synthesize c-di-GMP, suggesting a delayed swarmer to stalked cell differentiation in these cells. They proposed that upon pilus retraction, PilA enters the inner membrane and binds to and activates PleC kinase (Medico et al., 2020). This interaction with PilA might also play a role in PleC release from the cell pole and its switch to kinase activity.

### PAS domains allow modulation of enzymatic activity in response to specific signals

In this study we investigated how deletion of PleC’s PAS domains affects its function *in vivo*. Deletion of PleC PAS domains prevents PodJ from localizing PleC and inhibiting PleC kinase activity. This diffuse version of PleC exhibited increased PleD phosphorylation and WT levels of DivK dephosphorylation *in vivo* (Figure 3B and 6B). Mutant cells that have diffuse PleC(11PAS) as their only source of PleC retained their ability to grow pili and rotate their flagellum, suggesting that new pole development is not impaired in this mutant (Figure 4A, 4B, 6B, 6C, 6D, 6E). Together these results suggest that signal integration through PleC’s PAS domains primarily lead to modulations in kinase and not phosphatase activity.

By removing the PAS domains, PleC is now unable to properly integrate cellular cues, which results in an increase of PleD∼P relative to WT cells. While a burst of PleD phosphorylation by PleC initiates swarmer to stalked cell differentiation, the increased PleD∼P in *pleC(ΔPAS)* cells caused premature CtrA degradation (Figure 5A and 5B). We showed that this rapid CtrA degradation led to *pleC(ΔPAS)* cells spending less time as swarmer cells, negatively affecting their ability to swarm on plates (Figure 3C, 5C and 5D). Thus, PleC PAS domain interaction with PodJ allows proper timing of PleD phosphorylation, first by physically separating PleC from PleD in stalked and pre-divisional cells, and by inhibition of kinase activity prior the start of swarmer to stalked cell differentiation.

We note that inhibition of kinase activity by PodJ might not be the sole function of the PleC PAS domains. DivK dephosphorylation by PleC located at the pole opposite of DivJ kinase activity on DivK has previously been suggested to contribute to an asymmetric distribution of DivK∼P (Matroule et al., 2004; Tsokos et al., 2011). The spatial separation of DivK∼P and DivK, together with the asymmetric distribution of c-di-GMP (Christen et al., 2010) and the CtrA∼P gradient (Lasker et al., 2020), allow for a robust integrated system that ensures proper differentiated cell fates of *Caulobacter* daughter cells. This redundancy seems to tolerate the diffuse localization of PleC(*Δ*PAS), but proper PleC localization might become essential for cell cycle progression in cells where CtrA∼P or c-di-GMP spatial distribution perturbed.

PAS domains are found to integrate multiple cellular signals in different organisms (Möglich et al., 2009). Currently, PodJ is the only known interaction partner that directly binds to PleC’s PAS domains. However, it was shown that the enzymatic state of *Caulobacter’s* histidine kinase, CckA, depends on integration of multiple signals through its two PAS domains, PAS-A and PAS-B. Specifically, high CckA density, sensed by PAS-A self-association and the presence of unbound DivL, sensed by PAS-B, promotes CckA kinase activity, while high c-di-GMP concentration and the presence of the DivL-DivK∼P complex, sensed through PAS-B, inhibits CckA kinase and promotes phosphatase activity (Mann et al., 2016; Mann & Shapiro, 2018). It is still unclear whether PleC PAS domains respond to factors other than PodJ. The hypermotility observed in *pleC(ΔPAS)* cells was unexpected and could be the result of aberrant, mis-localized kinase activity on unknown targets. The effect of the PAS domains on PleC’s function reveals that cells can utilize a single histidine kinase to integrate cellular signals in a tailored response, allowing both phosphorylation and dephosphorylation of diverse targets, modulated by the presence or absence of context dependent binding partners.

### Temporal regulation of cell differentiation as a possible means for adaptation

The timing of swarmer to stalked cell differentiation, which was inferred by observation of either the onset of chromosome replication or the appearance of the stalk pole-specific protein DivJ, occurred on average, approximately six minutes sooner for *pleC(ΔPAS)* cells compared to WT cells (Figure 5C and 5D). While this difference in swarmer cell duration seems like a short amount time, we anticipate that this small time difference could result in major biological consequences for *Caulobacter* growing in the wild. *Caulobacter’s* di-morphic lifestyle allows it to disperse and colonize new habitats where it can form biofilms, which offer the advantage of increased tolerance to physical and chemical stressors. This di-morphic lifestyle enables *Caulobacter* to thrive in low nutrient environments such as lakes and streams (Poindexter, 1964). A swarmer duration that is six minutes shorter translates to a shorter distance that cells are able to disperse.

*Caulobacter* in the wild experiences environmental changes such as shifts in pH, temperature, and availability of nutrients. The ability of the species to utilize both lifestyles to its advantage hinges on *Caulobacter’s* ability to integrate signals from the environment into the decision of whether or not to differentiate. A previous study found that under conditions of carbon starvation, *Caulobacter* undergoes the initial morphological changes associated with stalked pole morphogenesis but stalls chromosome replication, not fully committing to the stalked cell fate (Britos et al., 2011). A recent publication showed that the activity of the histidine kinase, DivJ, is modulated by changes in the physical properties of the IDR-induced biomolecular condensate of its interaction partner, SpmX. This study specifically showed that the SpmX IDR responds to changes in ATP levels, whereby low levels of ATP promote condensate formation, further stimulating DivJ kinase and preventing cell division defects associated with low DivJ kinase activity (Saurabh et al., 2022). PleC PAS domains might play an important role in regulating PleC kinase activity under changing environmental conditions. As PleC PAS domains interact with PodJ’s IDR, it is tempting to consider that changes in the physical properties of PodJ’s IDR in response to environmental changes might also play a role in modulating PleC kinase activity, and ultimately regulate the timing of cell differentiation under specific environmental conditions. The periplasmic Cache domain of PleC might also play a role in regulating PleC enzymatic activity. We showed that deleting the periplasmic Cache domain led to reduced polar localization of PleC, suggesting that PleC interaction with a factor in the periplasm likely contributes to its polar localization (Figure 2B). Cache domains are sensory domains that typically bind to proteins or small molecules resulting in protein conformational changes that ultimately impact enzymatic activity (Upadhyay et al., 2016). Changes in environmental conditions could result in Cache domain-mediated regulation of PleC kinase activity and therefore timing of cell differentiation.

The timing of cell differentiation is a critical factor in unicellular and multicellular development, that when mis-regulated can have severe biological consequences. Our analysis of the dynamic nature of PleC kinase and phosphatase activities, mediated by the interaction of the PleC PAS domains with the PodJ polar scaffold, enables the temporal regulation of cell differentiation. These results highlight the complexity of two component systems and how bacterial kinases have evolved to integrate multiple signals into enzymatic activities functioning in time and space that result in distinct biological outcomes.

## Acknowledgements

We thank Sofia Calayag for help with developing the single cell tracking protocol. We thank Seth Childers and Edward de Koning for helpful discussions as well as all members of the L.S. lab. This work is supported by an NSFGRFP fellowship to T.C. and an NIH grant to L.S. (R35-GM118071). L.S. is a Chan Zuckerberg Biohub Investigator.

## Author Contributions

T.C. performed the experiments with help from S.S. and M.P.; T.C. and L.S. designed experiments and analyzed data with help from S.S.; T.C. and L.S. prepared the figures and wrote the manuscript.

## Resource Availability

All plasmids and bacterial strains generated by this study are available upon request to Lucy Shapiro or Saumya Saurabh without restriction.

All code generated by this study is available through Zenodo: https://doi.org/10.5281/zenodo.7796548

## Materials and Methods

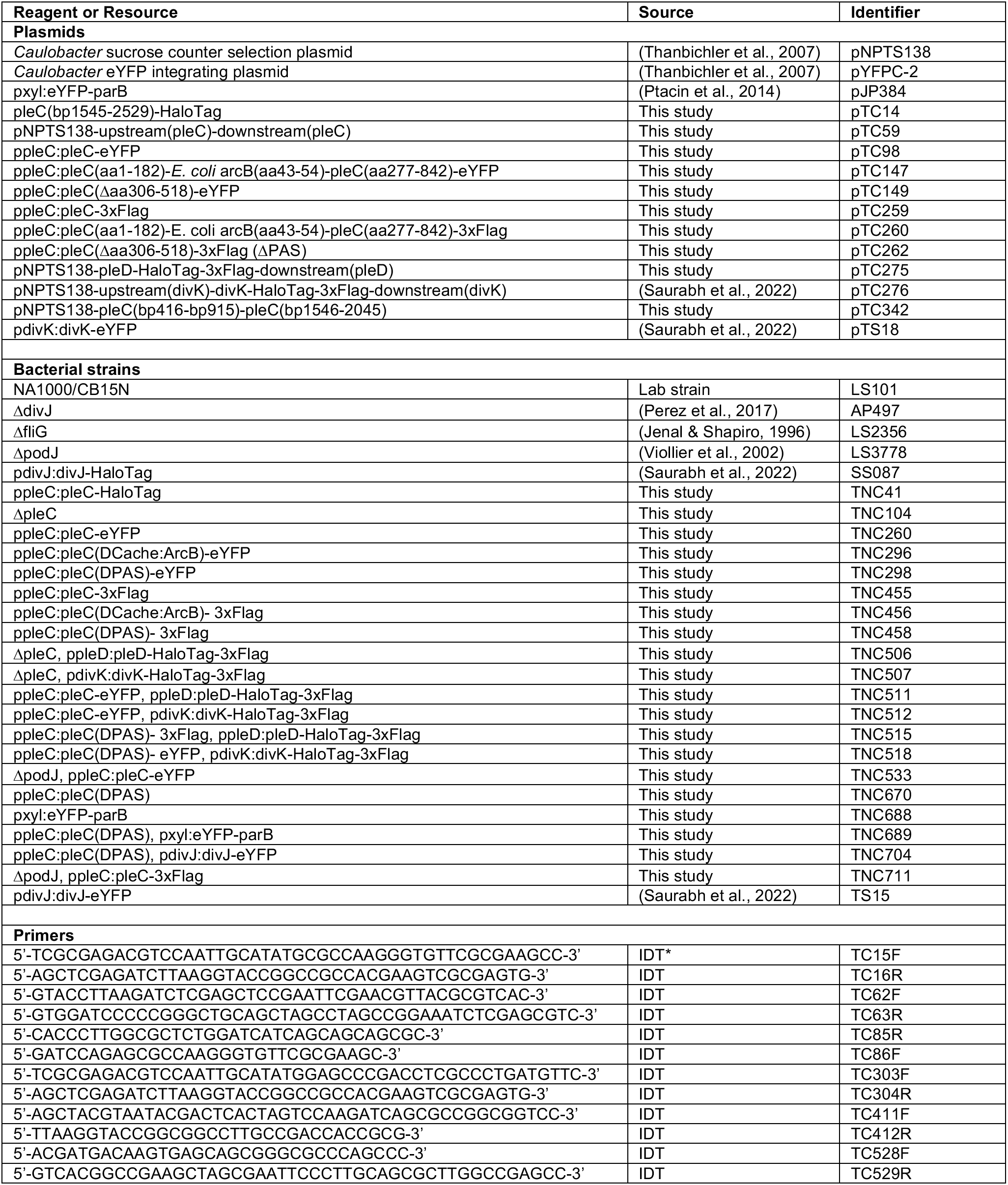
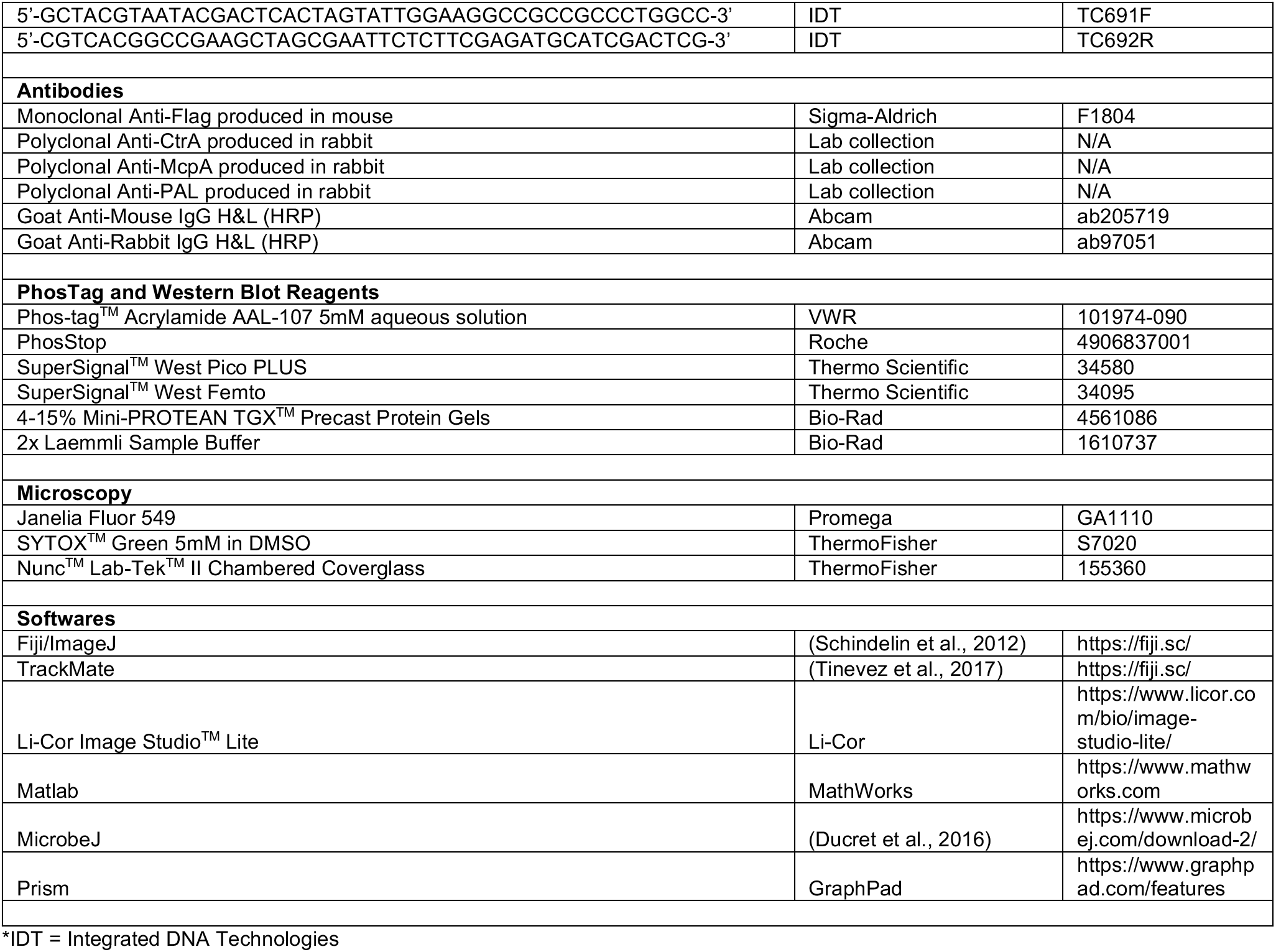

### Growth conditions

Unless otherwise stated, *Caulobacter* cells were grown from frozen stocks in PYE under aerating conditions at 30°C. Overnight PYE cultures were diluted accordingly into M2G or PYE in order to reach the desired OD600 the following day.

### Strain construction

All strains are derivatives of LS101, a lab stock of NA1000/CB15N. The strain TNC41 was made by electroporation of pTC14 into LS101. The plasmid pTC14 was constructed by Gibson assembly of pYFPC-2 digested with NdeI and KpnI and PCR amplification of PleC(bp1545-2529) from NA1000 genomic DNA with primers TC15F and TC16R. The resulting plasmid was digested with NheI and SacI, HaloTag amplified with primers TC62F and TC63R was inserted by Gibson assembly. TNC104 and TNC372 were made by electroporation of pTC59 into LS101 and LS3778 respectively, followed by sucrose counter selection. The plasmid pTC59 was constructed by Gibson assembly of pNPTS138 digested with SpeI and EcoRI and PCR amplifications from NA1000 genomic DNA with (TC190F and TC191R) and (TC192F and TC193R). TNC260, TNC296, TNC298, TNC455, TNC456 and TNC458 were made by electroporation of pTC98, pTC147, pTC149, pTC259, pTC60 and pTC262 into TNC104 respectively. The plasmid pTC98 was constructed by Gibson assembly of pYFPC-2 digested with NdeI and KpnI and PCR amplification from NA1000 genomic DNA with primers TC303F and TC304R. The plasmid pTC147 was constructed by Synbio (pTC98 was digested with NdeI and BmgBI and a synthesized DNA sequence containing ppleC:pleC(aa1-182)-*E. coli* ArcB(aa43-54)-pleC(aa277-393) was inserted). The plasmid pTC149 was constructed by Gibson assembly of pYFPC-2 digested with NdeI and KpnI and PCR amplifications from NA1000 genomic DNA with primers (TC303F and TC85R) and (TC86F and TC16R). The plasmid pTC259 was constructed by Synbio (pTC98 was digested with SacI and NheI and a synthesized DNA sequence encoding for 3xFlag was inserted). The plasmids pTC260 and pTC262 were constructed by Gibson assembly of pTC259 digested with NdeI and KpnI and PCR amplified regions from pTC147 and pTC149 with primers TC303F and TC304R. TNC506 and TNC507 were made by electroporation of pTC275 and pTC276 into TNC104 respectively, followed by sucrose counter selection. The plasmid pTC275 was constructed by Gibson assembly of pNPTS138 digested with SpeI and EcoRI with PCR amplifications from NA1000 genomic DNA with primers (TC411F and TC412R) and (TC528F and TC529R) and a DNA sequence synthesized by IDT encoding 3xFlag with overlapping sequences. TNC511 and TNC512 were made by electroporation of pTC98 into TNC506 and TNC507 respectively. TNC515 and TNC518 were made by electroporation of pTC149 into TNC506 and TNC507 respectively. TNC533 and TNC711 were made by electroporation of pTC98 and pTC259 into TNC372 respectively. TNC670 was made by electroporation of pTC342 into LS101 followed by sucrose counter selection. The plasmid pTC342 was constructed by Gibson assembly of pNTPS138 digested with SpeI and EcoRI and PCR amplifications from NA1000 genomic DNA with primers (TC691F and TC85R) and (TC86F and TC692R). TNC688 and TNC689 were made by electroporation of pJP384 into LS101 and TNC670 respectively. TNC704 was made by electroporation of pTS18 into TNC670.

### PhosTag

Cells were grown overnight in M2G to an OD600 of 0.3-0.5. Samples were normalized to an equivalent of 1mL of cells at OD600 of 0.4, and cell pellets were flash frozen in liquid nitrogen and stored at -80°C. Samples were lysed at room temperature for 5 minutes in 75μL of 10mM Tris-HCl (pH=7), 4% SDS, 2U/mL DNase, supplemented with PhosStop phosphatase inhibitor cocktail tablet, and then placed directly on ice. Samples were then centrifuged for 5 minutes at 13,000RPM and 12μL of the supernatant was then added to 12μL of Bio-Rad 2x Laemmli Sample Buffer with 5% φ-mercaptoethanol. From these samples, 20μL was loaded onto PhosTag gels. PhosTag poly-acrylamide gels were made with a final concentration of 100μM ZnCl_2_ as described by the Wako PhosTag guidebook. Phosphorylated and unphosphorylated bands were separated by gel electrophoresis at 4°C. Gels were washed three times in Towbin buffer containing 10mM EDTA for 10 minutes at room temperature followed by one wash without EDTA. Protein was transferred onto PVDF membranes by semi-dry transfer for 4 hours followed by standard Western Blot antibody probing procedures.

### Western Blot

Cells grown overnight in M2G were normalized to an equivalent of 1mL of cells at an OD600 of 0.4 and pellets were either flash frozen in liquid nitrogen and stored at -80°C or immediately lysed. To lyse, cells were resuspended in 40μL dH_2_O and combined with 40μL of Bio-Rad 2x Laemmli Sample Buffer with 5% φ-mercaptoethanol and boiled at 95°C for 10 minutes. Samples were loaded onto precast Bio-Rad 4-15% gradient poly-acrylamide gels followed by gel electrophoresis. Protein was transferred to PVDF membranes by semi-dry transfer for 2 hours. Sigma monoclonal anti-Flag M2 antibody (1:4,000 dilution) or Rabbit polyclonal anti-serra recognizing CtrA (1:10,000 dilution), McpA (1:20,000) or PAL (1:50,000 dilution) were used for immunoblots, followed by Abcam goat-anti-mouse HRP (1:10,000 dilution) or Abcam goat-anti-rabbit HRP (1:10,000 dilution). SuperSignal West Pico PLUS or SuperSignal West Femto Chemiluminescent Substrate from Thermo Scientific was used to detect HRP. Density measurements were calculated using Li-Cor Image Studio Lite. For CtrA synchrony signal density measurements, WT and *pleC(ΔPAS)* samples were run on the same gel.

### Synchrony

Cells were grown overnight in M2G to an OD600 of 0.4-0.5, 1L for large-scale or 40mL for small-scale synchronies. Cells were pelleted and washed with cold M2. Cells were resuspended in 1:1 M2 to Percoll and swarmer cells were separated from stalked and pre-divisional cells by density centrifugation (large scale: one hour at 6,000RPM, small scale: 20 minutes at 11,000RPM) at 4°C. The higher band consisting of stalks and pre-divisional cells was aspirated and swarmer cells were washed in cold M2 before being released into M2G and grown at 30°C. At appropriate time points cells were removed and normalized to an OD600 of 0.4 and flash frozen in liquid nitrogen and stored at -80°C.

### Imaging

Unless otherwise stated, all cell imaging was done on M2G 1.5% agarose pads on a light-emitting diode-based (Lumencor, SpectraX) multicolor epifluorescence microscope consisting of a Leica Dmi8 stand equipped with an immersion oil phase contrast objective [100×, HC PL APO, 1.4 numerical aperture (NA)] and an EMCCD camera (Hamamatsu, C9100 02 CL). Cells were grown overnight in M2G to an OD600 of 0.3-0.4. For generating line profiles, MicrobeJ was used to divide cells into either 26 or 50 bins along the longitudinal axis and the average fluorescence intensity of each bin was used to generate fluorescence profile plots. The protocol for imaging ϕCbK attachment was adapted from Hinz et al (Hinz et al., 2003). Briefly, Sytox Green was added to 1mL of ϕCbK to a final concentration of 25μM and allowed to incubate overnight at 4°C. Cells were mixed 1:1 with fluorescently labeled phage and imaged.

### Time-lapse imaging

For time-lapse imaging, swarmer cells isolated from a small-scale synchrony were either prepared for imaging immediately or allowed to grow to the appropriate stage of the cell cycle prior to imaging. For HaloTag imaging, cells were incubated in 2nM JF549 dye in M2G for 10 minutes at room temperature and washed 3 times with M2G prior to imaging. Images were taken at 10-minute time intervals. For PleC-HaloTag, cells were labeled immediately after synchrony. For DivJ-HaloTag, swarmer cells were allowed to progress to stalked cells for 40 minutes prior to labeling. For imaging DivJ-eYFP and eYFP-ParB cells, isolated swarmer cells were grown in M2G or an M2G with 0.01% xylose for 1 hour to allow cells to progress to the pre-divisional stage. Agarose pads made with M2G or M2G with 0.01% xylose were cut in half in order to image WT and *pleC(ΔPAS)* cells simultaneously. The microscope was programmed to take images at 5-minute intervals and move the stage to take images of both halves of the agarose pad. The genotypes associated with raw images were blinded and Δt intervals were acquired manually.

### Single cell tracking

For single cell tracking, glass bottom wells were treated with 1M KOH for 30 minutes and then washed 3 times with dH_2_O. Swarmer cells isolated from a small-scale synchrony were resuspended in 50-200μL of cold M2. Prior to imaging, 10μL of cells in cold M2 were added to 190μL of room temperature M2G with 20% glycerol and allowed to assimilate for at least 3 minutes. Images were taken at a focal plane above the coverslip in order to not capture cells that had settled to the bottom of the well. Images were captured at 52ms intervals. The Fiji plugin, TrackMate, was used to track individual cells and calculate speeds.

### Plate assays

For all plate assays, images were taken with a Bio-Rad ChemiDoc MP Imager. For swarm assays, single colonies were poked into PYE 0.3% agar plates and allowed to grow at 30°C for 3 days prior to imaging. Swarm areas were calculated using Matlab and averaged over replicate plates. For plaque assays, 400μL of mid-log cells were combined with 4mL melted PYE with 0.3% agar and poured on top of regular PYE plates (1.5% agar) and cooled to room temperature. 1.5μL of 10x serial dilutions of ϕCbK phage were spotted onto plates. Plates were incubated at room temperature overnight and images were taken the next day. To assay for cell growth, 1.5μL of 1:1 mixtures of mid-log cells with 10x serial dilutions of ϕCbK were spotted onto PYE agar plates. Plates were incubated at room temperature overnight, and images were taken the following day.

## Supplemental Materials

**Figure S1.**
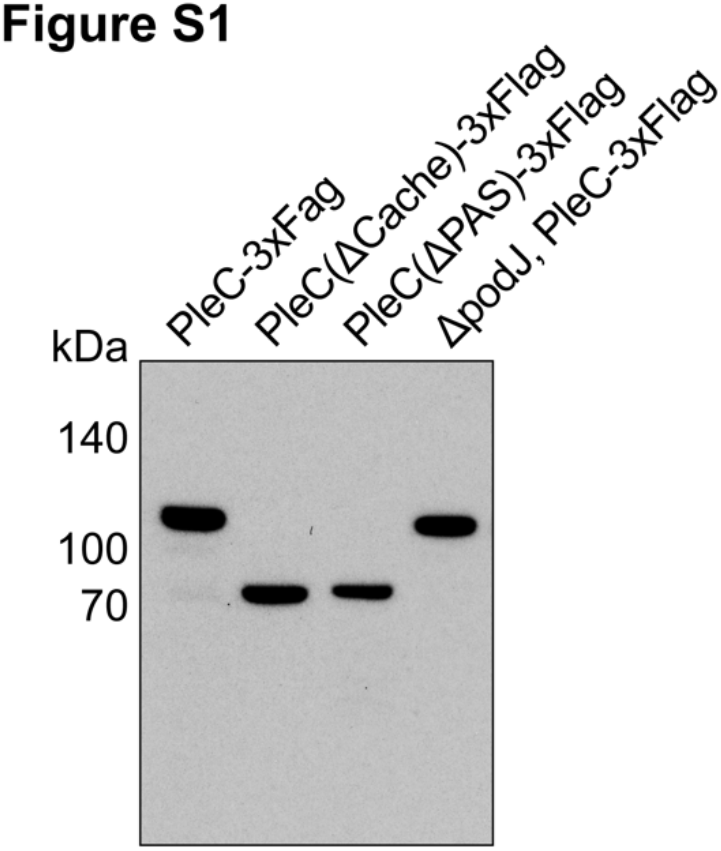
Western blot of PleC domain deletions C-terminally fused to 3xFlag and expressed as the sole copy of PleC from the endogenous locus and WT PleC-3xFlag in *ΔpodJ* probed with an anti-Flag antibody.

**Figure S2.**
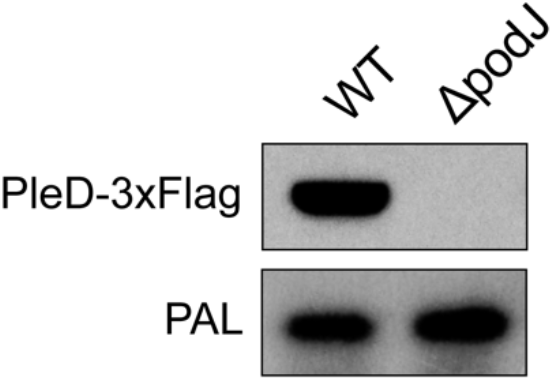
Western blot of WT and *ΔpodJ* cells expressing PleD-HaloTag-3xFlag at the *pleD* locus as the sole copy of PleD probed with an anti-Flag antibody and an anti-PAL antibody.

